# Tree species shape ectomycorrhizal fungal traits and community composition through litter quality and resultant soil properties

**DOI:** 10.1101/2025.10.06.680628

**Authors:** Castro David, Lyu Qian, Gil-Martínez Marta, Vesterdal Lars, Kjøller Rasmus

## Abstract

Ectomycorrhizal (ECM) fungi are key components of the nitrogen (N) and carbon (C) cycles in temperate and boreal forests soils. However, it is still unclear how differences in soil properties driven by litter quality shape ECM fungal diversity and composition. To test this, we used a unique common garden experiment in Denmark which included three common European tree species, lime, beech, and Norway spruce, each associated with contrasting litter quality from fast-to slow-turnover respectively. We found that soil physicochemical properties varied significantly among tree species, with spruce soils having higher soil organic matter (SOM), phosphorus (P), inorganic N and lower pH, than beech and lime. A total of 1,248 ECM root tips were selected for Sanger sequencing and individually assessed for the presence of hyphae and/or rhizomorphs protruding from the fungal mantle. Sequencing revealed 133 fungal amplicon sequence variants (ASV’s), clustered in 86 fungal operational taxonomic units (OTU’s), from which 61 corresponded to ECM fungi. Compared to spruce, lime and beech supported a higher ECM fungal richness and diversity in lime and beech compared to spruce. The ECM fungal community composition differed significantly between broadleaved species and spruce, with *Russula* dominating in lime and beech. Regarding fungal traits, in addition to the direct observations of hyphae and rhizomorphs, species were also scored based on literature for exploration types and hyphae hydrophobicity. Tree species had a specific association with ECM fungal exploration types, with broadleaved species favoring contact and short-distance types and spruce associating with long-distance types. The results suggest that tree species via foliar litter quality have a direct impact on soil properties and indirect to the ECM fungal community, where broadleaved tree species with nutrient-rich and rapid decomposing litter support a more diverse ECM community with limited soil exploration capabilities, while spruce supports an ECM fungal community more adapted to soil exploration.

## 1 Introduction

Tree species differ in their nutrient economy (Phillips et al., 2013) and C cycling dynamics (Vesterdal et al., 2012, Mayer et al., 2020) and with an anticipated increase in climatic and anthropogenic perturbations, an important question is how different tree species influence C and N cycling and the soil organisms that govern these processes. Notably, the effect on forest C and N cycling is pronounced, primarily due to significant tree species differences in litter quality and turnover rates (Vesterdal et al., 2008). Carbon turnover and N retention/mobilisation are mediated by the soil biota (Peng et al., 2022a, Zheng et al., 2022a), which plays a vital role in litter decomposition, N mobilization and retention, and C storage (Wardle and Lindahl, 2014). One of the key fungal players that directly links trees with soil are the symbiotic root colonizing mycorrhizal fungi (Brundrett, 2009). In temperate and boreal forests, the most abundant symbiosis occurs with ectomycorrhizal (ECM) fungi.

The tree nutrient economy is driven by the combination of inherent soil properties, symbiotic root-fungus feedback and tree litter quality. Litter quality can be characterized by the combination of nutrient concentrations, stoichiometry (C:N:P ratios), and specific lignocellulose chemistry (Zak et al., 2019). Indeed, litter quality is the main driver of decomposition rates globally (Cornwell et al., 2008). To better understand the impact of host species on the ECM fungal community, it is essential to study ECM fungal communities among different hosts growing in the same soil and with similar environmental conditions. Prior studies have investigated the effects of tree species on ECM fungi by comparing tree species growing in different site conditions (Rosinger et al., 2018, Otsing et al., 2021, Khokon et al., 2023). However, this approach can lead to confusion, as the effects of tree species might be confounded with pre-existing environmental differences among sites.

Ectomycorrhizal fungi exchange C, and thereby energy, directly from living trees, for N and P obtained via exploration and decomposition of soil organic matter (SOM) (Lindahl and Tunlid, 2015). There are large differences between ECM fungi in inherent mycelial production, extracellular enzymatic activity and mycelium hydrophobicity – traits of importance for exploration, acquisition and mobilization of nutrients from the soil (Kjøller, 2006, Kohler et al., 2015, Lindahl and Tunlid, 2015, Romero-Olivares et al., 2021, Unestam and Sun, 1995, Lilleskov et al., 2011). In general, species with hydrophilic ECM root tip mantles and short distance mycelia are thought to thrive under conditions with ample and labile available nutrients while the presence of hydrophobic mantle tissues and rhizomorphs is a strategy to prevent leakage of solutes taken up from organic nutrient patches at longer distance from the root (Lilleskov et al., 2011). Ectomycorrhizal fungal communities therefore respond to, or adapt to different soil fertility regimes, particularly N-status (Lilleskov et al., 2002, Cox et al., 2010, Kjøller et al., 2012). The tree hosts are to some degree thought to select the current ECM mycobiome from the available meta-community by allocating more or less C belowground to specific fungi (Law et al., 2022). Ectomycorrhizal traits for nutrient acquisition (*e*.*g*., soil exploration or hyphae hydrophobicity) are usually scored by combining observed species lists with trait databases (*e*.*g*., Determination of Ectomycorrhizae database (DEEMY) (Rambold and Agerer, 1997), FUNGuild (Nguyen et al., 2016), FungalTraits (Põlme et al., 2021), and Romero-Olivares et al. (2021)). However, nutrient acquisition traits may also be scored from in situ material as originally described by Agerer (2001) and operationalized for many samples by López-García et al. (2018). Overall, the environment, the available meta-community and host specificity thus all drive the local assemblage of ECM fungi at specific locations (Ishida et al., 2007).

The aim of this study was to investigate the influence of tree species with inherent differences in litter quality, *i*.*e*., Norway spruce (*Picea abies* (L.) Karst.), European beech (*Fagus sylvatica* L.) and small-leaved lime (*Tilia cordata* L.), on their associated ECM communities. Ectomycorrhizal communities are here understood as both the taxonomic units present on the root tips and as aggregated response traits. The three tree species can be ranked by increasing leaf litter quality, *i*.*e*., N, P and base cations from spruce over beech and to lime. Spruce produces recalcitrant, slowly decomposing litter which results in pronounced accumulation of forest floors and low soil pH. Lime, at the other end, produces high quality and fast decomposing litter, higher soil pH and sparse forest floor build-up (Vesterdal et al., 2013). To study these interactions, we took advantage of a unique common garden tree species experiment in Denmark established in 1973. At five sites, tree species were planted in adjacent 0.25 ha plots (Vesterdal et al., 2008, Vesterdal et al., 2012), locally on the same parent material and shared land use history. The biogeochemistry, soil biota and enzyme stoichiometry has been studied previously and provide reference and metadata for our study (Vesterdal et al., 2008, Heděnec et al., 2020, Zheng et al., 2022a, Zheng et al., 2022b). Fungal species were identified by Sanger sequencing single ECM root tips. Ectomycorrhizal root tips morphology was assessed by direct microscopy of sampled ECM root tips for presence of hyphae and rhizomorphs (López-García et al., 2018). In addition, fungal traits were recorded as exploration types and hyphae hydrophobicity by mapping database values on the species and genera detected.

We hypothesized that foliar litter quality would have a significant impact on ECM fungal community composition, diversity and their morphological soil exploratory traits. Specifically we hypothesize that 1) the root ECM fungal richness and diversity would reflect the stoichiometry associated with different litter quality inputs, and decrease as soil nutrient availability decreases, 2) the taxonomic composition would reflect the different soil properties derived from the tree host litter qualities *i*.*e*. a stronger response to tree species than for site location and 3) morphological traits, specifically mycelial exploration types (based on ECM root morphology and functional annotation) and hydrophobicity, are expected to change directionally along the gradient of species-specific foliar litter quality. We expected that these in situ measured traits would be mirrored by mapping exploration types to the taxonomic composition *i*.*e*. contact and short distance exploration types and hydrophilic ECM hyphae are likely to dominate in soils with high nutrient availability, whereas long distance and hydrophobic ECM hyphae are expected to prevail under low-nutrient soils.

## 2 Methods

### 2.1 Study site

Three common temperate tree species were studied, the two broadleaved species, beech (*Fagus sylvatica* L.) and lime (*Tilia cordata* L.), and the coniferous Norway spruce (*Picea abies* (L.) Karst.). These species were selected from the common garden established across Denmark in 1973 (Vesterdal et al., 2008). Three sites, Odsherred, Vallø and Viemose were selected for this study given their similarities in former land-use, soil characteristics and geographical proximity (Vesterdal et al., 2008) (Fig. 1A; 1B). Each species was planted in monoculture plots of about 0.25 ha. The annual average precipitation ranged from 580 to 890 mm, while the annual average temperature ranged from 7.5ºC to 8.4ºC (Heděnec et al., 2020). The elevations of the sites ranged from 13 to 68 meters above sea level. The soil type present in all locations were Stagnic Luvisols. A more detailed description of the sites can be found in Vesterdal et al. (2008) and Heděnec et al. (2020).

**Figure 1.**
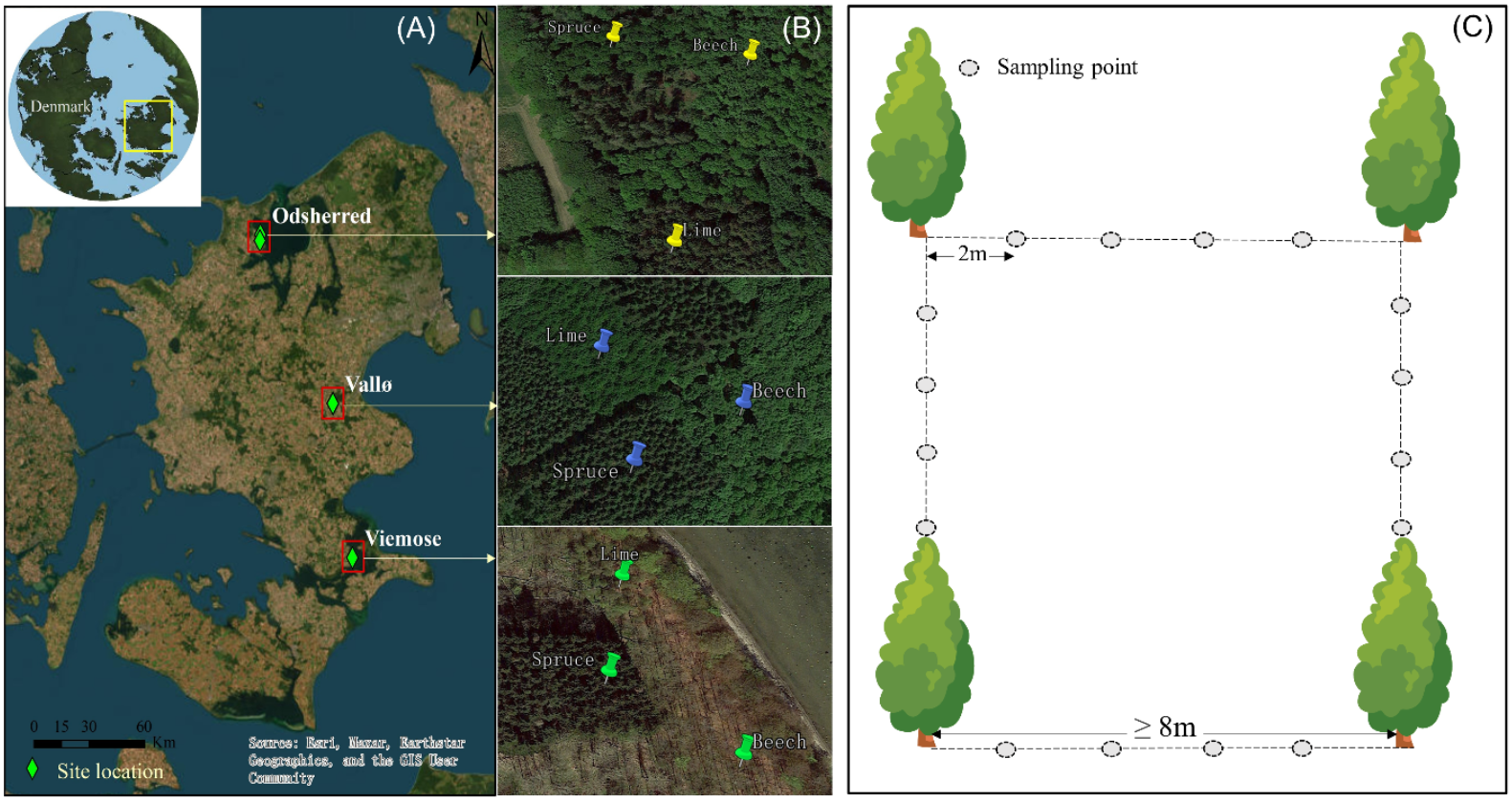
Location map of the three sites (A), plots at individual sites (B) and sampling design within the individual plots (C).

### 2.2 Foliar litter characterization

Previous studies on the common gardens found that lime has the highest foliar litter quality compared to other ECM tree species, including beech and spruce (Vesterdal et al., 2012, Peng et al., 2022b). As such, lime foliar litter presented higher N and P contents, as well as lower lignin:N ratio (Table 1). In addition, lime litter has higher contents of K, Ca and Mg (Table 1). On the other hand, spruce has lower contents of N, P and higher lignin:N ratio (Table 1). A detailed description of the foliar litter quality of these species can be found in Vesterdal et al. (2012) and Peng et al. (2022b).

**Table 1.** Foliar litter quality of the three tree species: lime, beech and spruce. Values correspond to mean values ± standard error from three sites together. Data from Vesterdal et al. (2012).

| Species | C<br>(mg·g<sup>-1</sup>) | N<br>(mg·g<sup>-1</sup>) | C:N | P<br>(mg·g<sup>-1</sup>) | Lignin:N | K<br>(mg·g<sup>-1</sup>) | Ca<br>(mg·g<sup>-1</sup>) | Mg<br>(mg·g<sup>-1</sup>) | Mn<br>(mg·g<sup>-1</sup>) | Fe<br>(mg·g<sup>-1</sup>) |
| --- | --- | --- | --- | --- | --- | --- | --- | --- | --- | --- |
| Lime | 486±4<sup>a</sup> | 17±0.6<sup>a</sup> | 33±3<sup>ab</sup> | 1.1±0.1<sup>a</sup> | 15±0.8<sup>b</sup> | 5.6±0.7<sup>a</sup> | 16±2<sup>a</sup> | 2.2±0.13<sup>a</sup> | 1147±133<sup>a</sup> | 137±9<sup>a</sup> |
| Beech | 475±9<sup>a</sup> | 13±1.8<sup>ab</sup> | 37±4<sup>a</sup> | 0.9±0.1<sup>ab</sup> | 23±2.3<sup>a</sup> | 3.2±0.1<sup>ab</sup> | 12±1<sup>ab</sup> | 1.4±0.04<sup>b</sup> | 1148±36<sup>a</sup> | 386±303<sup>a</sup> |
| Spruce | 467±11<sup>a</sup> | 11±0.7<sup>b</sup> | 32±4<sup>b</sup> | 0.8±0.1<sup>b</sup> | 20±0.7<sup>ab</sup> | 2.7±0.1<sup>b</sup> | 10±0.2<sup>b</sup> | 1.0±0.03<sup>c</sup> | 1156±67<sup>a</sup> | 90±6<sup>a</sup> |

### 2.3 Sampling design

In November 2022, four trees approximately 8 m apart forming the corners of a square plot were randomly selected within each plot (Fig. 1C). To assess the ECM fungal community, four soil cores were taken from the top 10 cm of the forest floor (*i*.*e*., any organic layer and the top mineral soil layer) with a 2 cm diameter soil corer at 2 m intervals between trees (Fig. 1C). An additional set of four soil cores were collected and pooled for soil physicochemical analysis. A total of 16 soil cores for ECM analysis and 4 composite soil samples for soil analysis were taken per plot per site, amounting to a total of 144 soil samples for ECM analysis and 36 for soil analysis. All samples were stored at 4ºC until analysis.

### 2.4 Root tip morphology

Roots from soil samples were collected and washed with cold tap water on a 0.4 mm sieve after which root fragments with ECM roots were collected. Ectomycorrhizal root tips were observed under a stereo microscope (Olympus SZX16, Olympus, Denmark), photographed and recorded for the presence of emanating hyphae and/or rhizomorphs (Gil-Martínez et al., 2021). In total, 1,248 ECM root tips were successfully selected and summarized into four categories, as described by López-García et al. (2018): 1) ECM root tips without emanating hyphae or rhizomorphs, 2) with hyphae (*i*.*e*., ECM root tips with emanating hyphae), 3) with rhizomorphs only and finally 4) with both (*i*.*e*., ECM root tips with both emanating hyphae and rhizomorphs).

### 2.5 DNA extraction and sequencing

Genomic DNA from the 1,248 ECM root tips was extracted using the Extract-N-Amp™ Plant PCR Kit (Sigma-Aldrich, St. Louis, Missouri, United States). Briefly, a single root tip was incubated in 10µl of extraction solution at 95ºC for 10 min and after adding 10µl dilution solution the samples were incubated further 10 min at 20ºC. The mix including the remaining ECM tip tissue was used directly as DNA template for amplification. Amplification of the fungal rRNA ITS2 region was performed using the 5x HOT FIREPol Blend Master Mix (Soils BioDyne, Tartu, Estonia) using the ITS1F (Gardes and Bruns, 1993) and ITS4 (White et al., 1990) to a final volume of 20µl. The samples were then amplified using 35 cycles of annealing at 55ºC for 35 s. Amplification success was confirmed by electrophoresis and amplifications containing two or more bands were removed. In total, 1,152 PCR products were sent for Sanger sequencing on a 3730xl DNA Analyzer platform (Macrogen Europe, the Netherlands) using their standard sequencing service.

### 2.6 Raw data preprocessing and functional annotation

Sequence preprocessing was performed using QIIME2 v2024.5 (Bolyen et al., 2019). Briefly, before importing into QIIME2, Sanger files were converted into FASTA format and compiled into a single FASTA file using a custom made script. The compiled FASTA file was then imported to QIIME2 using the q2-import tool. Sequence quality control was performed using the UNITE database v.10.0 (Kõljalg et al., 2020) as reference using the VSEARCH method, with an identity percentage and query alignment over 97%. Quality controlled sequences were then annotated using q2-feature-classifier (Bokulich et al., 2018) plugin in combination with the classify-sklearn method (Pedregosa et al., 2011) to assign taxonomic annotation. This method uses a machine learning classifier trained with the v10.0 database of UNITE. Annotated fungal taxa were then summarised by Taxon and TaxaID using a custom made python script.

In total, 1,152 sequences were annotated into 133 fungal amplicon sequence variants (ASV’s) and clustered into 86 fungal operational taxonomic units (OTU’s). To obtain only the ECM fungi from the dataset, functional annotation was performed using FUNGuild v1.1 for guild annotation of all fungi, keeping only the fungi containing “Ectomycorrhizal” within their guild. In total, 307 sequences clustered in 61 fungal OTU’s taxa were kept as putative ECM fungi. Additionally, FungalTraits (Põlme et al., 2021) and DEEMY (https://www.deemy.de) databases were used to obtain fungal traits, such as exploration type, as described in Agerer (2001) and hyphal hydrophobicity (Unestam and Sun, 1995). Published references were used to annotate the fungal taxa when information was missing. See supplementary material for more details.

### 2.7 Soil physicochemical analyses

Composited soil samples were aliquoted for different physicochemical analyses. For gravimetrical soil water content fresh soil samples were passed through a 2 mm sieve and weighed, then air-drying at 80ºC until constant weight. Soil organic matter content was determined by loss on ignition at 550ºC for 6 h from the dry composited soil samples. Soil pH was measured in a 1:2.5 soil-water suspension after shaking for 30 min. Total soil C and N were estimated by dry combustion (Dumas method) in a Leco CNS 2000 analyser (Leco, St. Joseph, Michigan, United States). Soil ammonium, nitrate and phosphate were extracted using a 1:2.5 soil-water suspension. Ammonium was determined by salicylic acid-hypochlorite colorimetry, nitrate was measured by copper plated cadmium column-diazotization coupling colorimetry, and phosphate was measured by ascorbate molybdenum blue colorimetry. Determination of ammonium, nitrate, and phosphate were done using an AutoAnalyzer 500 (Seal Analytical, Norderstedt, Germany). Ammonium, nitrate and phosphate are expressed in mg per g of C.

### 2.8 Statistical analysis

All statistical analyses were conducted in R, version 4.4.0 (R Core Team 2024). As a single plot was established inside each monoculture plot per site, the site was included in the analysis as a random factor and no Species x Site interactions could be tested. Data normality was tested using Shapiro test and transformed using Tukey’s Ladder of Powers if normality assumption was not achieved. Soil properties were tested using multivariate analysis of variance (MANOVA) and visualized with a principal component analysis (PCA) plot using fviz_pca_biplot function from the factoextra package (Kassambara and Mundt, 2020). If significant differences were found in the MANOVA, one-way analysis of variance (ANOVA) and Tukey’s Honest Significant Differences (Tukey’s HSD) as post-hoc analysis were performed. To test the differences in ECM root tips morphology between tree species, one-way ANOVA and Tukey’s HSD were performed.

To assess the effect of the host species on the ECM fungal community diversity; richness, Shannon diversity index and Pielou’s evenness were performed with the vegan package (Oksanen et al., 2025). β-diversity, assessed as ECM community dissimilarities, was performed using vegdist from vegan with Bray-Curtis dissimilarity indexing. Statistical differences in richness, Shannon diversity index and Pielou’s evenness were assessed using one-way ANOVA and Tukey’s HSD. Differences in β-diversity between tree species was assessed using permutational multivariate analysis of variance (PerMANOVA) using the vegan package and Kruskal-Wallis when differences were significant.

Differences in ECM fungal traits, *i*.*e*. exploration types and hydrophobicity were tested using one-way ANOVA and Tukey’s HSD if significant differences were found. All plots were made with ggplot2 (Wickham, 2016). All script used in this study are available in a Github repository accompanying this work (https://github.com/davcastrom/EuroTreeECM).

## 3 Results

### 3.1 Soil properties

Significant physicochemical differences between spruce and broadleaved tree species (*i*.*e*., lime and beech) were found (MANOVA; *p-value* < 0.001; Fig. 2). Soils from spruce stands had significantly higher concentrations of total N, C and SOM, but the lowest pH (Table 2). On the other hand, the two broadleaved species did not differ significantly in their soil properties (Table 2). No significant differences in ammonium, nitrate and phosphate per gram of C were found between spruce and broadleaved species (Table 2).

**Table 2.** Soil chemical properties. Values correspond to mean values ± standard error from three sites. SOM = soil organic matter.

| Species | Ammonium (mg·gC<sup>-1</sup>) | C (mg·g<sup>-1</sup>) | C:N | Total N (mg·g<sup>-1</sup>) | Nitrate (mg·gC<sup>-1</sup>) | pH | Phosphate (mg·gC<sup>-1</sup>) | SOM (%) | Water Content (%) |
| --- | --- | --- | --- | --- | --- | --- | --- | --- | --- |
| Lime | 1.4±0.7<sup>a1</sup> | 49±7<sup>b</sup> | 26±6<sup>a</sup> | 2.2±0.8<sup>b</sup> | 1.9±1.3<sup>a</sup> | 5.5±0.2<sup>a</sup> | 0.6±0.2<sup>a</sup> | 9.9±1.3<sup>b</sup> | 34±4<sup>a</sup> |
| Beech | 1.0±0.4<sup>a</sup> | 60±19<sup>b</sup> | 43±31<sup>a</sup> | 2.7±2.2<sup>b</sup> | 1.6±1.3<sup>a</sup> | 5.5±0.5<sup>a</sup> | 0.5±0.2<sup>a</sup> | 13±4<sup>b</sup> | 40±12<sup>a</sup> |
| Spruce | 2.4±2.2<sup>a</sup> | 171±66<sup>a</sup> | 26±2.2<sup>a</sup> | 7.1±2.4<sup>a</sup> | 2.7±2.3<sup>a</sup> | 4.5±0.0<sup>b</sup> | 0.5±0.4<sup>a</sup> | 29±9<sup>a</sup> | 43±14<sup>a</sup> |

<sup>1</sup>Lowercase letters correspond to significant differences between tree species from Tukey's HSD test.

**Figure 2.**
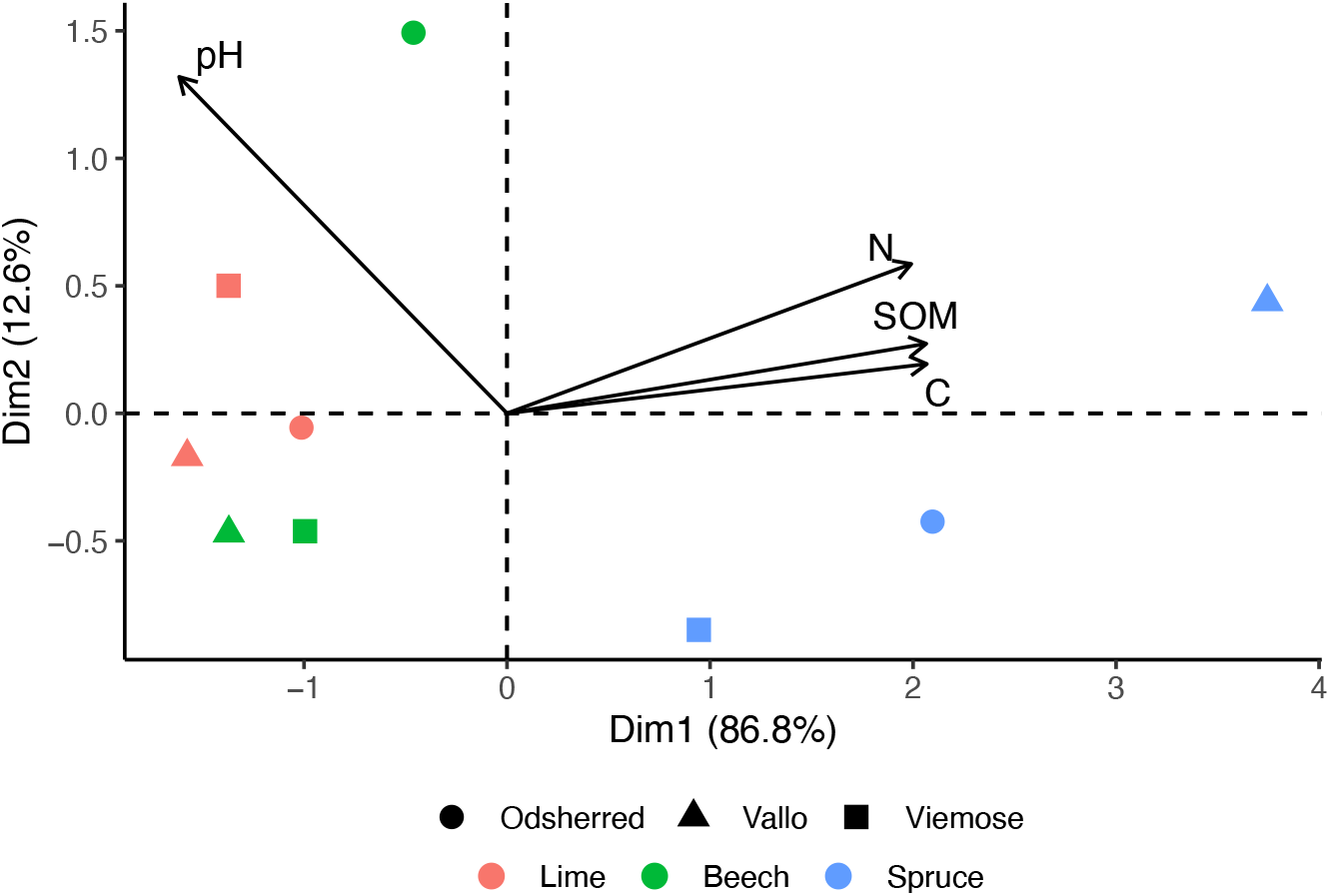
Principal component analysis (PCA) biplot of soil properties of soil samples from three different temperate tree species. Vectors correspond to the significantly different (*p-value* < 0.05) soil physicochemical properties. Dots correspond to each single composited soil sample, colored according to tree species and symbols according to site.

### 3.2 ECM fungal community composition and diversity

From the 1,248 sequenced root tips, 133 fungal amplicon sequence variants (ASV’s), clustered in 86 fungal operational taxonomic units (OTU’s) were identified. Functional annotation using FUNGuild showed that lime had the largest relative abundance of ECM fungi, followed by beech (Fig. 3). Fine root tips from spruce, on the other hand, showed a higher relative abundance of other biotrophic fungi (*i*.*e*., lichenized, ericoid mycorrhizal and endophytic), compared to broadleaved trees (Fig. 3). In addition, spruce had the largest relative abundance of ECM fungi with unidentified guilds (*i*.*e*., ECM fungi annotated up to class without guild annotation in FUNGuild), *e*.*g*. Boletales, Chantharellales, and Russulales (Supplementary material).

**Figure 3.**
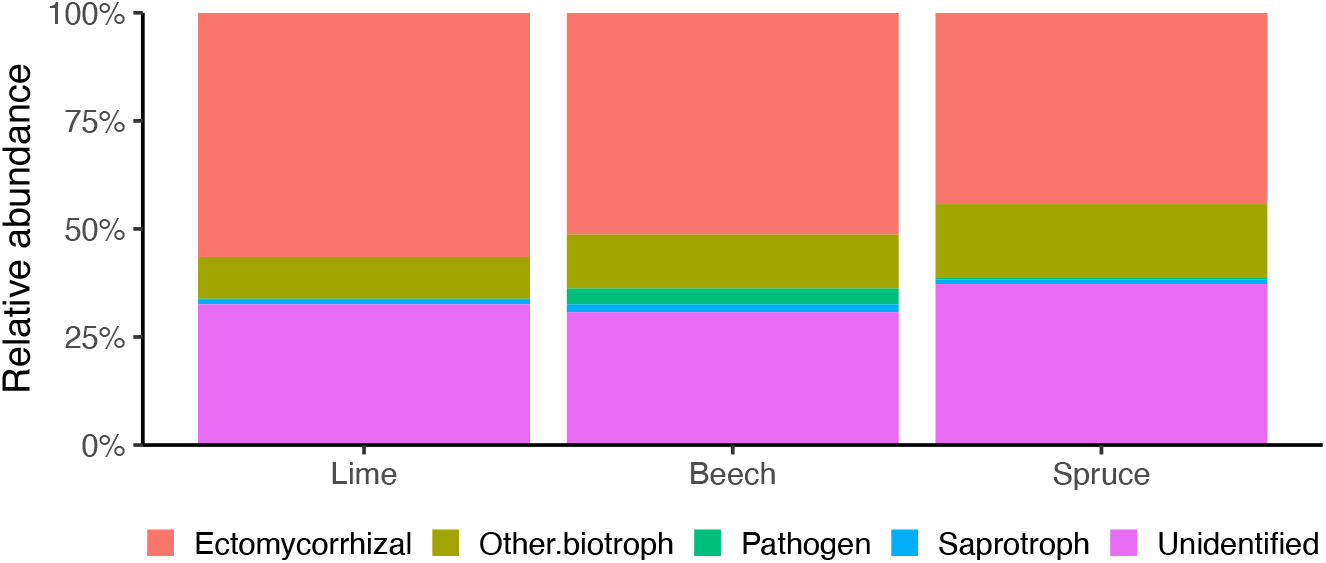
Whole root-associated fungal community from root sampled from three different temperate tree species. Relative abundance of functional guilds colored by guild.

Community composition was evaluated with non-metric multidimensional scaling using Bray-Curtis dissimilarity index. Goodness of fit (r^2^) suggests that the site explains 26% of the ECM community variations, while the tree species had a significant impact on the ECM community variation, explaining 52%. Furthermore, the dissimilarity test suggested a significant difference in ECM fungal genera between tree species, with spruce having a higher dissimilarity index when compared with the broadleaved species than when comparing beech with lime (PerMANOVA; *p-value* < 0.001; Fig. S1). This difference was mostly explained by the clear dominance of *Russula* in both lime and beech compared to spruce. In addition to *Russula* species, unidentified *Thelephoraceae* species and *Cortinarius* were the second and third most abundant ECM genera in lime, while in beech *Lactarius* and *Tomentella* were the second and third most abundant ECM genera (Table 3). Furthermore, lime and beech shared six ECM genera that were absent in spruce, such as *Cortinarius, Elaphomyces, Laccaria, Piloderma, Sebacina* and *Thelephora* (Table 3). Finally, *Hydnum, Inocybe, Pachyphlodes* and *Tuber* were unique in lime, and *Entomola, Genea, Humaria, Hydnotrya*, and *Leotia* were unique in beech (Table 3). In contrast to beech and lime, spruce ECM fungal community was dominated by unidentified *Thelephoraceae* species, followed by *Amphinema* and *Russula* (Table 3). Four genera were unique in spruce (*i*.*e*., *Amphinema, Imleria, Tylopilus* and *Tylospora*). Seventeen genera present in lime and/or beech were on the other hand absent on spruce (Table 3).

**Table 3.**
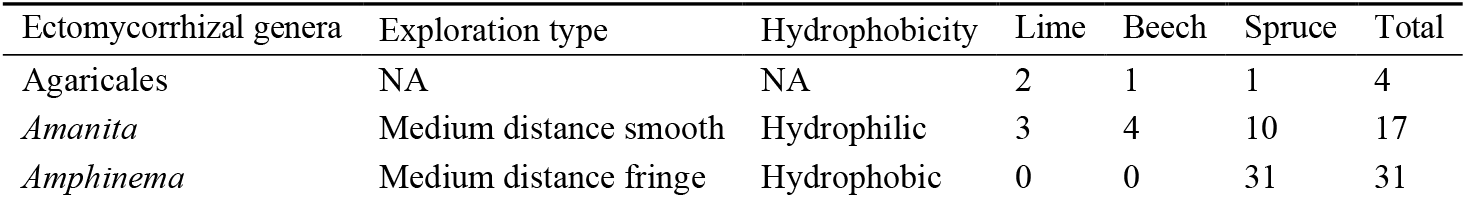
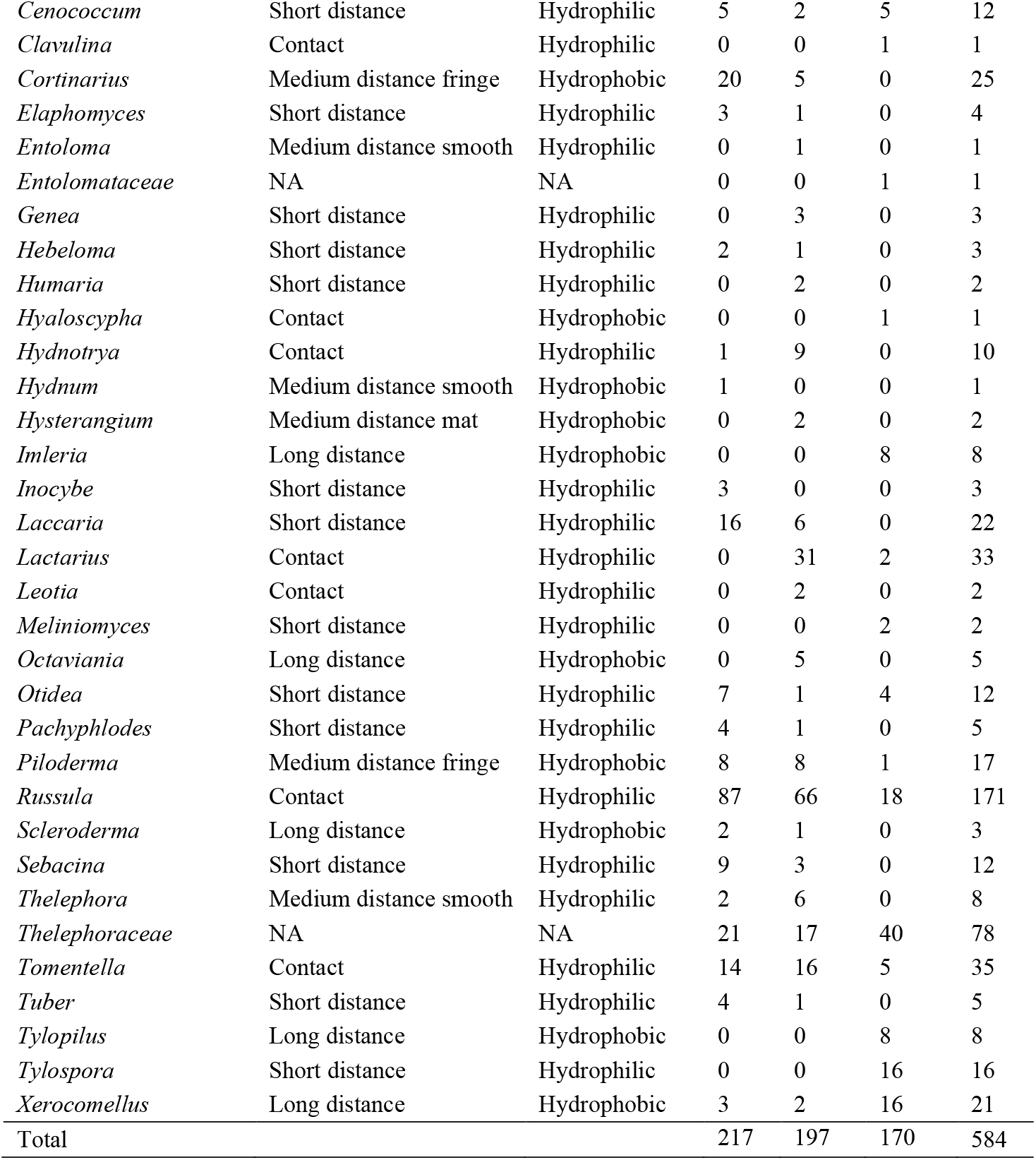
Ectomycorrhizal genera in three temperate tree species sorted by name. Values correspond to the sum of annotated ECM from n = 12 samples per tree species. A detailed version of this data can be found in the supplementary material.

| Ectomycorrhizal genera | Exploration type | Hydrophobicity | Lime | Beech | Spruce | Total |
| --- | --- | --- | --- | --- | --- | --- |
| Agaricales | NA | NA | 2 | 1 | 1 | 4 |
| *Amanita* | Medium distance smooth | Hydrophilic | 3 | 4 | 10 | 17 |
| *Amphinema* | Medium distance fringe | Hydrophobic | 0 | 0 | 31 | 31 |
| *Cenococcum* | Short distance | Hydrophilic | 5 | 2 | 5 | 12 |
| *Clavulina* | Contact | Hydrophilic | 0 | 0 | 1 | 1 |
| *Cortinarius* | Medium distance fringe | Hydrophobic | 20 | 5 | 0 | 25 |
| *Elaphomyces* | Short distance | Hydrophilic | 3 | 1 | 0 | 4 |
| *Entoloma* | Medium distance smooth | Hydrophilic | 0 | 1 | 0 | 1 |
| *Entolomataceae* | NA | NA | 0 | 0 | 1 | 1 |
| *Genea* | Short distance | Hydrophilic | 0 | 3 | 0 | 3 |
| *Hebeloma* | Short distance | Hydrophilic | 2 | 1 | 0 | 3 |
| *Humaria* | Short distance | Hydrophilic | 0 | 2 | 0 | 2 |
| *Hyaloscypha* | Contact | Hydrophobic | 0 | 0 | 1 | 1 |
| *Hydnotrya* | Contact | Hydrophilic | 1 | 9 | 0 | 10 |
| *Hydnum* | Medium distance smooth | Hydrophobic | 1 | 0 | 0 | 1 |
| *Hysterangium* | Medium distance mat | Hydrophobic | 0 | 2 | 0 | 2 |
| *Imleria* | Long distance | Hydrophobic | 0 | 0 | 8 | 8 |
| *Inocybe* | Short distance | Hydrophilic | 3 | 0 | 0 | 3 |
| *Laccaria* | Short distance | Hydrophilic | 16 | 6 | 0 | 22 |
| *Lactarius* | Contact | Hydrophilic | 0 | 31 | 2 | 33 |
| *Leotia* | Contact | Hydrophilic | 0 | 2 | 0 | 2 |
| *Meliniomyces* | Short distance | Hydrophilic | 0 | 0 | 2 | 2 |
| *Octaviania* | Long distance | Hydrophobic | 0 | 5 | 0 | 5 |
| *Otidea* | Short distance | Hydrophilic | 7 | 1 | 4 | 12 |
| *Pachyphlodes* | Short distance | Hydrophilic | 4 | 1 | 0 | 5 |
| *Piloderma* | Medium distance fringe | Hydrophobic | 8 | 8 | 1 | 17 |
| *Russula* | Contact | Hydrophilic | 87 | 66 | 18 | 171 |
| *Scleroderma* | Long distance | Hydrophobic | 2 | 1 | 0 | 3 |
| *Sebacina* | Short distance | Hydrophilic | 9 | 3 | 0 | 12 |
| *Thelephora* | Medium distance smooth | Hydrophilic | 2 | 6 | 0 | 8 |
| *Thelephoraceae* | NA | NA | 21 | 17 | 40 | 78 |
| *Tomentella* | Contact | Hydrophilic | 14 | 16 | 5 | 35 |
| *Tuber* | Short distance | Hydrophilic | 4 | 1 | 0 | 5 |
| *Tylopilus* | Long distance | Hydrophobic | 0 | 0 | 8 | 8 |
| *Tylospora* | Short distance | Hydrophilic | 0 | 0 | 16 | 16 |
| *Xerocomellus* | Long distance | Hydrophobic | 3 | 2 | 16 | 21 |
| Total |  |  | 217 | 197 | 170 | 584 |

These differences in ECM fungal composition were reflected in significant differences in richness, with beech having significantly higher species richness of ECM fungi than spruce (Fig. 4A). These differences in ECM species richness were reflected in significant differences in Shannon diversity between beech and spruce (Fig. 4B). There were no significant differences in ECM evenness among the three tree species (Fig. 4C).

**Figure 4.**
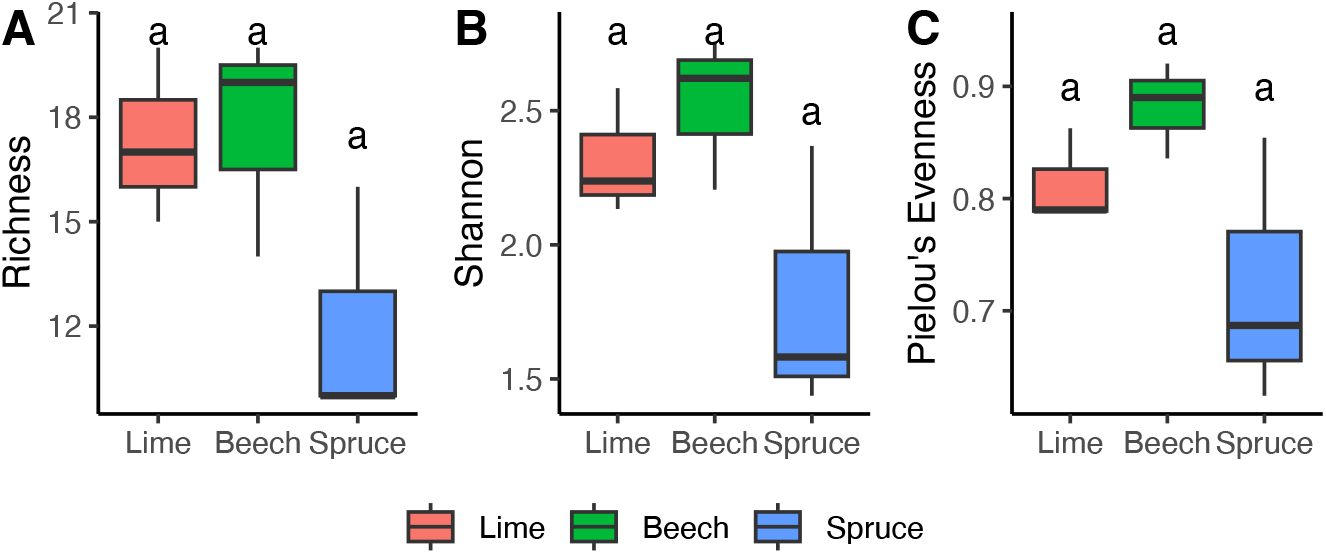
Boxplots of ECM fungal species richness and diversity in stands of lime, beech and spruce. Lowercase letters above the boxplot correspond to Tukey’s HSD when *p-value* < 0.05 from one-way ANOVA test of differences between tree species was found. N = 3.

### 3.3 ECM fungal traits: exploration type and hydrophobicity

In total, 398, 392, and 458 root tips from lime, beech and spruce, respectively, were observed under the stereo microscope for root tip morphology annotation. From the 1,248 selected root tips, 501 were successfully annotated as ECM and were also annotated for exploration type and hydrophobicity (Table 4). Lime root tips accounted for the highest numbers of root tips without hyphae or rhizomorphs, as well as only hyphae accounting for 50% of all the root tips (Table 4). Beech had the largest number of only rhizomorphs, and hyphae and rhizomorph root tips, followed by spruce (Table 4). Interestingly, both beech and spruce accounted for 31% of root tips containing rhizomorphs.

**Table 4.** Summary of fungal traits per tree species in all sites expressed as number of root tips successfully sequenced and annotated in each category. Relative abundance in % by tree species within parentheses.

|  | Lime | Beech | Spruce | Total |
| --- | --- | --- | --- | --- |
| **Morphology (*in situ*)** |  |  |  |  |
| No hyphae or rhizomorphs | 74 (38)<sup>a1</sup> | 54 (30)<sup>a</sup> | 43 (34)<sup>a</sup> | 171 |
| Only hyphae | 97 (50)<sup>a</sup> | 68 (38)<sup>ab</sup> | 45 (35)<sup>b</sup> | 210 |
| Only rhizomorphs | 6 (3)<sup>b</sup> | 24 (13)<sup>a</sup> | 12 (9)<sup>ab</sup> | 42 |
| Hyphae and rhizomorphs | 17 (9)<sup>a</sup> | 33 (18)<sup>a</sup> | 28 (22)<sup>a</sup> | 78 |
| Total | 194 (100) | 179 (100) | 128 (100) | 501 |
| **Exploration type** |  |  |  |  |
| Contact | 102 (53)<sup>a</sup> | 124 (69)<sup>a</sup> | 27 (21)<sup>b</sup> | 253 |
| Short distance | 53 (27)<sup>a</sup> | 21 (12)<sup>a</sup> | 27 (21)<sup>a</sup> | 101 |
| Medium distance fringe | 28 (14)<sup>a</sup> | 13 (7)<sup>a</sup> | 32 (25)<sup>a</sup> | 73 |
| Medium distance mat | 0 (0)<sup>a</sup> | 2 (1)<sup>a</sup> | 0 (0)<sup>a</sup> | 2 |
| Medium distance smooth | 6 (3)<sup>a</sup> | 11 (6)<sup>a</sup> | 10 (8)<sup>a</sup> | 27 |
| Long distance | 5 (3)<sup>b</sup> | 8 (4)<sup>b</sup> | 32 (25)<sup>a</sup> | 45 |
| Total | 194 (100) | 179 (100) | 128 (100) | 501 |
| **Hydrophobicity** |  |  |  |  |
| Hydrophilic | 160 (82)<sup>a</sup> | 156 (87)<sup>a</sup> | 63 (49)<sup>b</sup> | 379 |
| Hydrophobic | 34 (18)<sup>a</sup> | 23 (13)<sup>a</sup> | 65 (51)<sup>a</sup> | 122 |
| Total | 194 (100) | 179 (100) | 128 (100) | 501 |

<sup>1</sup>Lowercase letters correspond to significant differences between tree species from Tukey's HSD test.

When assessing the sequenced data for fungal traits, broadleaved tree species showed a prevalence of contact and short distance exploration types, covering both more than 75% of the community (Table 4). Norway spruce showed a significant preference for long distance exploration type ECM, covering about 25% of the community, followed by the contact and short distance exploration types (Table 4). Medium distance mat was the lowest for all tree species (Table 4). Regarding hyphal hydrophobicity, lime and beech presented a significant prevalence of hydrophilic hyphal fungi, while spruce did not show a significant difference (Table 4).

## 4 Discussion

In this study, we present the influence of tree species-specific foliar litter quality inputs on the soil properties and the cascading effect on the ECM fungal community. As expected, we found large effects of the inherent tree-species litter qualities on soil properties which in turn had a significant impact on the ECM community, diversity and ECM fungal traits. Broadleaved tree species (lime and beech) had richer soils than spruce. The higher soil nutrient availability, found in lime and beech soils, related to a richer and more diverse ECM community than for spruce, which confirmed our first hypothesis. In concordance with our second hypothesis, we found that the ECM community composition was influenced more by tree host species (52%) than by location (26%). Finally, in support of our third hypothesis, a significantly higher prevalence of long-distance exploration types and hydrophobic ECM fungi were found in spruce compared to the two broadleaved species. A novel aspect of our study was that the prevalence of rhizomorphs and long-distance exploration types was found both from mapping database trait values onto our taxonomic composition as well as by direct observations of root tips extracted from the field.

The impact of tree foliar litter on the soil properties is well known. For instance, tree species with high lignin content produce a thicker forest floor. This difference has also been established between arbuscular mycorrhizal (AM) and ECM tree species, with the latter producing more recalcitrant foliar litter. Previous studies at our common garden locations site have found that lime produces high quality foliar litter, behaving more closely to the litter quality of the AM tree species ash and maple, than other ECM tree species like beech or spruce (Peng et al., 2022a). Beech foliar litter is known to be more recalcitrant than other broadleaved ECM species and particular that of lime, having higher C:N and lignin content (Yang et al., 2019). This is consistent with the foliar litter data from Vesterdal et al. (2012) (Table 1), where beech had significantly higher C:N and lignin:N ratio than lime. However, despite significant differences in litter quality between lime and beech, we did not find significant differences in soil properties between the two broadleaved tree species however both were significant different from spruce (Fig. 2 and Table 2).

Richness was markedly higher for lime and beech compared with spruce. This is consistent with previous results across Europe. For instance, a meta-analysis study reported higher fungal richness in beech compared to spruce or pine (Rosinger et al., 2018). Similarly, Heděnec et al. (2023) found significant differences in fungal richness between lime and beech compared with spruce, with the later having 11 and 12 less OTU’s, respectively. A hypothesis could be that the combination of both organic matter and more available nutrients below lime and beech compared with spruce allows for a wider number of species with different nutrient acquisition traits co-exist. Under spruce a narrower set of species adapted to mine out nutrients from the more recalcitrant organic matter are found. A recent study in a former *Betula pubescens* site in Norway has showed that the establishment of Norway spruce significantly reduced the fungal richness and diversity in the litter and humus layer (Danielsen et al., 2021). Bahnmann et al. (2018) found in a mixed forest stand in the Czech Republic that a significantly lower proportion of ECM fungi in spruce compared to beech. In our results, we also found lower ECM richness in spruce trees, which can be partially attributed to an acidified soil as a consequence of its recalcitrant litter (Vesterdal et al., 2012, Peng et al., 2022b). The soil under spruce may be thought of as a stressful environment and only stress-tolerant species may thrive. The abundance of species with hydrophobic mantles may be seen as a trait linked to surviving in a stressed environment (Castaño et al., 2023, Defrenne et al., 2019). All our three sites were before 1973 beech forests/plantations and our present day communities can be attributed to diverging ECM recruitment since then. An alternative hypothesis could therefore be that more species were capable to continue from the former beech forest to the new beech and lime plots. Finally, it should be noted that spruce did not on its own make it back to Denmark after the past glaciation (but was present in the area before). The lower richness under spruce may therefore also partly be attributed to fewer compatible fungi within the region.

Our data showed that ECM communities mainly clustered according to tree species rather than geography. Both soil properties and the host tree species have been recognized as drivers for the soil ECM fungal community. For instance, Correia et al. (2021) found that the land-use history, with higher nitrate and ammonia, and lower organic matter content, significantly altered the ECM richness in newly established beech forests. Similarly, large scale experiments have recognized soil properties, particularly soil pH, as a driving force for ECM selection (Tedersoo et al., 2020, Odriozola et al., 2023). By using a common garden, we reduced the effect of distinct founder communities and soil properties associated with different land-use histories. From the start in 1973, all trees were planted in soil with similar history in each site, *i*.*e*. former beech forest or former cropland. What has happened later has largely been determined because of the different tree species influence. After almost 50 years of the common garden establishment, any legacies of the original fungal community composition at each site (from original resistant propagules) are probably minimal.

The differences in community structure were accompanied by a significantly higher prevalence of long-distance exploration type and mycelial hydrophobicity in spruce than in lime or beech (Fig. 5) corroborated by the presence of rhizomorphs on the sampled root tips (Table 4). Similar results have been found when comparing *Picea asperata* and *Larix gmelinii* with *Quercus aquifolioides* and *Betula albosinensis*, where *P. asperata* had significantly higher mycelial length, accompanied by a higher prevalence of long-distance exploration type and hydrophobic mycelial fungi (Xie et al., 2024). Previous studies in soil hydrology have suggested that sites with poor litter quality, podzolization processes and low pH (Flores-Mangual et al., 2013, González-Sosa et al., 2024), as well as poor foliar litter water retention (Feng et al., 2025) increases water repellency in the soil (*i*.*e*., more hydrophobic soils). In addition, it has been proposed that hydrophobic fungi are prevalent in hydrophobic forest soils, having greater adherence to the soil and a greater nutrient, especially organic N, acquiring capability (Lilleskov et al., 2011). This suggests that hydrophobic fungi follow a “like-to-like” pattern with the soil properties, presumably as a consequence of their greater capacity to obtain nutrients from the soil aggregates compared to the more hydrophilic fungi. This is supported by the finding of Almeida et al. (2022) who found a direct link between hydrophobic fungi and the soil organic matter in Norway spruce soils. The higher prevalence of long distance and hydrophobic ECM fungi in the spruce plot is an indicative of the strong impact of the foliar litter quality in the soil properties and the subsequent response of the ECM community to those changes.

## Acknowledge

QL appreciates the financial support from the China Scholarship Council (grant numbers 202206910054). MG-M was supported by “Margarita Salas” grant funded by the Spanish Recovery, Transformation and Resilience Plan and NextGenerationEU. DC, RK and LV were all supported by the Danish Research Council (grant no 3103-00085B).

## Data

GitHub and long-term sequence storage facility.

## Author contribution

QL: Experimental design, lab work, Draft writing, Formal analysis DC: Formal analysis, writing and editing, programming MG-M: Experimental design, lab work LV: Project management, writing and editing RK: Project management, experimental design, supervising, writing and editing

## References

2024. R: A Language and Environment for Statistical Computing.

Agerer, R. 2001. Exploration types of ectomycorrhizae. Mycorrhiza, 11, 107–114.

Almeida, J. P., Rosenstock, N. P., Woche, S. K., Guggenberger, G. & Wallander, H. 2022. Nitrophobic ectomycorrhizal fungi are associated with enhanced hydrophobicity of soil organic matter in a Norway spruce forest. Biogeosciences, 19, 3713–3726.

Bahnmann, B., Mašínová, T., Halvorsen, R., Davey, M. L., SedláK, P., Tomšovský, M. & Baldrian, P. 2018. Effects of oak, beech and spruce on the distribution and community structure of fungi in litter and soils across a temperate forest. Soil Biology and Biochemistry, 119, 162–173.

Bokulich, N. A., Kaehler, B. D., Rideout, J. R., Dillon, M., Bolyen, E., Knight, R., Huttley, G. A. & Gregory Caporaso, J. 2018. Optimizing taxonomic classification of marker-gene amplicon sequences with QIIME 2’s q2-feature-classifier plugin. Microbiome, 6, 90.

Bolyen, E., Rideout, J. R., Dillon, M. R., Bokulich, N. A., Abnet, C. C., Al-Ghalith, G. A., Alexander, H., Alm, E. J., Arumugam, M., Asnicar, F., Bai, Y., Bisanz, J. E., Bittinger, K., Brejnrod, A., Brislawn, C. J., Brown, C. T., Callahan, B. J., Caraballo-Rodriguez, A. M., Chase, J., Cope, E. K., Da Silva, R., Diener, C., Dorrestein, P. C., Douglas, G. M., Durall, D. M., Duvallet, C., Edwardson, C. F., Ernst, M., Estaki, M., Fouquier, J., Gauglitz, J. M., Gibbons, S. M., Gibson, D. L., Gonzalez, A., Gorlick, K., Guo, J., Hillmann, B., Holmes, S., Holste, H., Huttenhower, C., Huttley, G. A., Janssen, S., Jarmusch, A. K., Jiang, L., Kaehler, B. D., Kang, K. B., Keefe, C. R., Keim, P., Kelley, S. T., Knights, D., Koester, I., Kosciolek, T., Kreps, J., Langille, M. G. I., Lee, J., Ley, R., Liu, Y. X., Loftfield, E., Lozupone, C., Maher, M., Marotz, C., Martin, B. D., Mcdonald, D., Mciver, L. J., Melnik, A. V., Metcalf, J. L., Morgan, S. C., Morton, J. T., Naimey, A. T., Navas-Molina, J. A., Nothias, L. F., Orchanian, S. B., Pearson, T., Peoples, S. L., Petras, D., Preuss, M. L., Pruesse, E., Rasmussen, L. B., Rivers, A., Robeson, M. S., 2Nd, Rosenthal, P., Segata, N., Shaffer, M., Shiffer, A., Sinha, R., Song, S. J., Spear, J. R., Swafford, A. D., Thompson, L. R., Torres, P. J., Trinh, P., Tripathi, A., Turnbaugh, P. J., Ul-Hasan, S., Van Der Hooft, J. J. J., Vargas, F., Vazquez-Baeza, Y., Vogtmann, E., Von Hippel, M., Walters, W., et al. 2019. Reproducible, interactive, scalable and extensible microbiome data science using QIIME 2. Nat Biotechnol, 37, 852–857.

Brundrett, M. C. 2009. Mycorrhizal associations and other means of nutrition of vascular plants: understanding the global diversity of host plants by resolving conflicting information and developing reliable means of diagnosis. Plant and Soil, 320, 37–77.

Castaño, C., Suarez-Vidal, E., Zas, R., Bonet, J. A., Oliva, J. & Sampedro, L. 2023. Ectomycorrhizal fungi with hydrophobic mycelia and rhizomorphs dominate in young pine trees surviving experimental drought stress. Soil Biology and Biochemistry, 178.

Cornwell, W. K., Cornelissen, J. H., Amatangelo, K., Dorrepaal, E., Eviner, V. T., Godoy, O., Hobbie, S. E., Hoorens, B., Kurokawa, H., Perez-Harguindeguy, N., Quested, H. M., Santiago, L. S., Wardle, D. A., Wright, I. J., Aerts, R., Allison, S. D., Van Bodegom, P., Brovkin, V., Chatain, A., Callaghan, T. V., Diaz, S., Garnier, E., Gurvich, D. E., Kazakou, E., Klein, J. A., Read, J., Reich, P. B., Soudzilovskaia, N. A., Vaieretti, M. V. & Westoby, M. 2008. Plant species traits are the predominant control on litter decomposition rates within biomes worldwide. Ecol Lett, 11, 1065–71.

Correia, M., Espelta, J. M., Morillo, J. A., Pino, J. & RodríGuez-Echeverría, S. 2021. Land-use history alters the diversity, community composition and interaction networks of ectomycorrhizal fungi in beech forests. Journal of Ecology, 109, 2856–2870.

Cox, F., Barsoum, N., Lilleskov, E. A. & Bidartondo, M. I. 2010. Nitrogen availability is a primary determinant of conifer mycorrhizas across complex environmental gradients. Ecol Lett, 13, 1103–13.

Danielsen, J. S., Morgado, L., Mundra, S., Nybakken, L., Davey, M. & Kauserud, H. 2021. Establishment of spruce plantations in native birch forests reduces soil fungal diversity. FEMS Microbiol Ecol, 97.

Defrenne, C. E., Philpott, T. J., Guichon, S. H. A., Roach, W. J., Pickles, B. J. & Simard, S. W. 2019. Shifts in Ectomycorrhizal Fungal Communities and Exploration Types Relate to the Environment and Fine-Root Traits Across Interior Douglas-Fir Forests of Western Canada. Front Plant Sci, 10, 643.

Feng, Y., Zhao, Y., Zhang, D., Zhang, B., Fan, X., Han, Z. & Zhao, L. 2025. Response of water retention characteristics in litters based on the leaf type. Hydrology Research, 56, 297–304.

Flores-Mangual, M. L., Lowery, B., Bockheim, J. G., Pagliari, P. H. & Scharenbroch, B. 2013. Hydrophobicity of Sparta Sand under Different Vegetation Types in the Lower Wisconsin River Valley. Soil Science Society of America Journal, 77, 1506–1516.

Gardes, M. & Bruns, T. D. 1993. ITS primers with enhanced specificity for basidiomycetes--application to the identification of mycorrhizae and rusts. Mol Ecol, 2, 113–8.

Gil-Martínez, M., López-García, Á., Domínguez, M. T., Kjøller, R., Navarro-Fernández, C. M., Rosendahl, S. & Marañón, T. 2021. Soil fungal diversity and functionality are driven by plant species used in phytoremediation. Soil Biology and Biochemistry, 153, 108102, publisher = Pergamon.

González-Sosa, M., González-Barrios, P., Bentancur, O. J. & Pérez-Bidegain, M. 2024. Differential effects on soil water repellency of Eucalyptus and Pinus plantations replacing natural pastures. Revista Brasileira de Ciência do Solo, 48.

Heděnec, P., Nilsson, L. O., Zheng, H., Gundersen, P., Schmidt, I. K., Rousk, J. & Vesterdal, L. 2020. Mycorrhizal association of common European tree species shapes biomass and metabolic activity of bacterial and fungal communities in soil. Soil Biology and Biochemistry, 149, 107933.

Heděnec, P., Zheng, H., Pessanha Siqueira, D., Lin, Q., Peng, Y., Kappel Schmidt, I., Guldberg Frøslev, T., Kjøller, R., Rousk, J. & Vesterdal, L. 2023. Tree species traits and mycorrhizal association shape soil microbial communities via litter quality and species mediated soil properties. Forest Ecology and Management, 527, 120608, publisher = Elsevier.

Ishida, T. A., Nara, K. & Hogetsu, T. 2007. Host effects on ectomycorrhizal fungal communities: insight from eight host species in mixed conifer-broadleaf forests. New Phytol, 174, 430–440.

Kassambara, A. & Mundt, F. 2020. Factoextra: Extract and Visualize the Results of Multivariate Data Analyses. 1.0.7 ed.

Khokon, A. M., Janz, D. & Polle, A. 2023. Ectomycorrhizal diversity, taxon-specific traits and root N uptake in temperate beech forests. New Phytol, 239, 739–751.

Kjøller, R. 2006. Disproportionate abundance between ectomycorrhizal root tips and their associated mycelia. FEMS Microbiol Ecol, 58, 214–24.

Kjøller, R., Nilsson, L. O., Hansen, K., Schmidt, I. K., Vesterdal, L. & Gundersen, P. 2012. Dramatic changes in ectomycorrhizal community composition, root tip abundance and mycelial production along a stand-scale nitrogen deposition gradient. New Phytol, 194, 278–286.

Kohler, A., Kuo, A., Nagy, L. G., Morin, E., Barry, K. W., Buscot, F., Canback, B., Choi, C., Cichocki, N., Clum, A., Colpaert, J., Copeland, A., Costa, M. D., Dore, J., Floudas, D., Gay, G., Girlanda, M., Henrissat, B., Herrmann, S., Hess, J., Hogberg, N., Johansson, T., Khouja, H. R., Labutti, K., Lahrmann, U., Levasseur, A., Lindquist, E. A., Lipzen, A., Marmeisse, R., Martino, E., Murat, C., Ngan, C. Y., Nehls, U., Plett, J. M., Pringle, A., Ohm, R. A., Perotto, S., Peter, M., Riley, R., Rineau, F., Ruytinx, J., Salamov, A., Shah, F., Sun, H., Tarkka, M., Tritt, A., Veneault-Fourrey, C., Zuccaro, A., MYCORRHIZAL GENOMICS Initiative, C., Tunlid, A., Grigoriev, I. V., Hibbett, D. S. & Martin, F. 2015. Convergent losses of decay mechanisms and rapid turnover of symbiosis genes in mycorrhizal mutualists. Nat Genet, 47, 410–5.

Kõljalg, U., Nilsson, H. R., Schigel, D., Tedersoo, L., Larsson, K. H., May, T. W., Taylor, A. F. S., Jeppesen, T. S., Froslev, T. G., Lindahl, B. D., Põldmaa, K., Saar, I., Suija, A., Savchenko, A., Yatsiuk, I., Adojaan, K., Ivanov, F., Piirmann, T., Pohonen, R., Zirk, A. & Abarenkov, K. 2020. The Taxon Hypothesis Paradigm-On the Unambiguous Detection and Communication of Taxa. Microorganisms, 8.

Law, S. R., Serrano, A. R., Daguerre, Y., Sundh, J., Schneider, A. N., Stangl, Z. R., Castro, D., Grabherr, M., Näsholm, T., Street, N. R. & Hurry, V. 2022. Metatranscriptomics captures dynamic shifts in mycorrhizal coordination in boreal forests. Proc Natl Acad Sci U S A, 119, e2118852119.

Lilleskov, E. A., Fahey, T. J., Horton, T. R. & Lovett, G. M. 2002. Belowground Ectomycorrhizal Fungal Community Change over a Nitrogen Deposition Gradient in Alaska. Ecology, 83, 104–115.

Lilleskov, E. A., Hobbie, E. A. & Horton, T. R. 2011. Conservation of ectomycorrhizal fungi: exploring the linkages between functional and taxonomic responses to anthropogenic N deposition. Fungal Ecology, 4, 174–183.

Lindahl, B. D. & Tunlid, A. 2015. Ectomycorrhizal fungi - potential organic matter decomposers, yet not saprotrophs. New Phytol, 205, 1443–1447.

López-García, Á., Gil-Martínez, M., Navarro-Fernández, C. M., Kjøller, R., Azcón-Aguilar, C., Domínguez, M. T. & Marañón, T. 2018. Functional diversity of ectomycorrhizal fungal communities is reduced by trace element contamination. Soil Biology and Biochemistry, 121, 202–211.

Mayer, M., Prescott, C. E., Abaker, W. E. A., Augusto, L., Cécillon, L., Ferreira, G. W. D., James, J., Jandl, R., Katzensteiner, K., Laclau, J.-P., Laganière, J., Nouvellon, Y., Paré, D., Stanturf, J. A., Vanguelova, E. I. & Vesterdal, L. 2020. Tamm Review: Influence of forest management activities on soil organic carbon stocks: A knowledge synthesis. Forest Ecology and Management, 466, 118127.

Nguyen, N. H., Song, Z., Bates, S. T., Branco, S., Tedersoo, L., Menke, J., Schilling, J. S. & Kennedy, P. G. 2016. FUNGuild: An open annotation tool for parsing fungal community datasets by ecological guild. Fungal Ecology, 20, 241–248.

Odriozola, I., Martinović, T., MašíNová, T., Bahnmann, B. D., Machac, A., Sedlák, P., Tomšovský, M. & Baldrian, P. 2023. The spatial patterns of community composition, their environmental drivers and their spatial scale dependence vary markedly between fungal ecological guilds. Global Ecology and Biogeography, 33, 173–188.

Oksanen, J., Simpson, G. L., Blanchet, F. G., Kindt, R., Legendre, P., Minchin, P. R., O’hara, R. B., Solymos, P., Stevens, M. H. H., Szoecs, E., Wagner, H., Barbour, M., Bedward, M., Bolker, B., Borcard, D., Carvalho, G., Chirico, M., De Caceres, M., Durand, S., Antoniazi Evangelista, H. B., Fitzjohn, R., Friendly, M., Furneaux, B., Hannigan, G., Hill, M. O., Lahti, L., Mcglinn, D., Ouellette, M.-H., Ribeiro Cunha, E., Smith, T., Stier, A., Ter Braak, C. J. F. & Weedon, J. 2025. vegan: Community Ecology Package. 2.6-10 ed.

Otsing, E., Anslan, S., Ambrosio, E., Koricheva, J. & Tedersoo, L. 2021. Tree Species Richness and Neighborhood Effects on Ectomycorrhizal Fungal Richness and Community Structure in Boreal Forest. Front Microbiol, 12, 567961.

Pedregosa, F., Varoquaux, G., Gramfort, A., Michel, V., Thirion, B., Grisel, O., Blondel, M., Prettenhofer, P., Weiss, R., Dubourg, V., Vanderplas, J., Passos, A., Cournapeau, D., Brucher, M., Perrot, M. & Duchesnay, É. 2011. Scikit-learn: Machine Learning in Python. Journal of Machine Learning Research, 12, 2825–2830.

Peng, Y., Holmstrup, M., Kappel Schmidt, I., Ruggiero Bachega, L., Schelfhout, S., Zheng, H., Heděnec, P., Yue, K. & Vesterdal, L. 2022a. Tree species identity is the predominant modulator of the effects of soil fauna on leaf litter decomposition. Forest Ecology and Management, 520, 120396.

Peng, Y., Holmstrup, M., Schmidt, I. K., De Schrijver, A., Schelfhout, S., Heděnec, P., Zheng, H., Bachega, L. R., Yue, K. & Vesterdal, L. 2022b. Litter quality, mycorrhizal association, and soil properties regulate effects of tree species on the soil fauna community. Geoderma, 407.

Phillips, R. P., Brzostek, E. & Midgley, M. G. 2013. The mycorrhizal-associated nutrient economy: a new framework for predicting carbon-nutrient couplings in temperate forests. New Phytol, 199, 41–51.

Põlme, S., Abarenkov, K., Henrik Nilsson, R., Lindahl, B. D., Clemmensen, K. E., Kauserud, H., Nguyen, N., Kjøller, R., Bates, S. T., Baldrian, P., Frøslev, T. G., Adojaan, K., Vizzini, A., Suija, A., Pfister, D., Baral, H.-O., Järv, H., Madrid, H., Nordén, J., Liu, J.-K., Pawlowska, J., Põldmaa, K., Pärtel, K., Runnel, K., Hansen, K., Larsson, K.-H., Hyde, K. D., Sandoval-Denis, M., Smith, M. E., Toome-Heller, M., Wijayawardene, N. N., Menolli, N., Reynolds, N. K., Drenkhan, R., Maharachchikumbura, S. S. N., Gibertoni, T. B., Læssøe, T., Davis, W., Tokarev, Y., Corrales, A., Soares, A. M., Agan, A., Machado, A. R., Argüelles-Moyao, A., Detheridge, A., De Meiras-Ottoni, A., Verbeken, A., Dutta, A. K., Cui, B.-K., Pradeep, C. K., Marín, C., Stanton, D., Gohar, D., Wanasinghe, D. N., Otsing, E., Aslani, F., Griffith, G. W., Lumbsch, T. H., Grossart, H.-P., Masigol, H., Timling, I., Hiiesalu, I., Oja, J., Kupagme, J. Y., Geml, J., Alvarez-Manjarrez, J., Ilves, K., Loit, K., Adamson, K., Nara, K., Küngas, K., Rojas-Jimenez, K., Bitenieks, K., Irinyi, L., Nagy, L. G., Soonvald, L., Zhou, L.-W., Wagner, L., Aime, M. C., Öpik, M., Mujica, M. I., Metsoja, M., Ryberg, M., Vasar, M., Murata, M., Nelsen, M. P., Cleary, M., Samarakoon, M. C., Doilom, M., Bahram, M., Hagh-Doust, N., Dulya, O., Johnston, P., Kohout, P., Chen, Q., Tian, Q., Nandi, R., Amiri, R., Perera, R. H., Dos Santos Chikowski, R., et al. 2021. FungalTraits: a user-friendly traits database of fungi and fungus-like stramenopiles. Fungal Diversity, 105, 1–16.

Rambold, G. & Agerer, R. 1997. DEEMY - the concept of a characterization and determination system for ectomycorrhizae. Mycorrhiza, 7, 113–116.

Romero-Olivares, A. L., Morrison, E. W., Pringle, A. & Frey, S. D. 2021. Linking Genes to Traits in Fungi. Microb Ecol, 82, 145–155.

Rosinger, C., Sandén, H., Matthews, B., Mayer, M. & Godbold, D. L. 2018. Patterns in Ectomycorrhizal Diversity, Community Composition, and Exploration Types in European Beech, Pine, and Spruce Forests. Forests, 9.

Tedersoo, L., Anslan, S., Bahram, M., Drenkhan, R., Pritsch, K., Buegger, F., Padari, A., Hagh-Doust, N., Mikryukov, V., Gohar, D., Amiri, R., Hiiesalu, I., Lutter, R., Rosenvald, R., Rahn, E., Adamson, K., Drenkhan, T., Tullus, H., Jurimaa, K., Sibul, I., Otsing, E., Polme, S., Metslaid, M., Loit, K., Agan, A., Puusepp, R., Varik, I., Koljalg, U. & Abarenkov, K. 2020. Regional-Scale In-Depth Analysis of Soil Fungal Diversity Reveals Strong pH and Plant Species Effects in Northern Europe. Front Microbiol, 11, 1953.

Unestam, T. & Sun, Y.-P. 1995. Extramatrical structures of hydrophobic and hydrophilic ectomycorrhizal fungi. Mycorrhiza, 5, 301–311.

Vesterdal, L., Clarke, N., Sigurdsson, B. D. & Gundersen, P. 2013. Do tree species influence soil carbon stocks in temperate and boreal forests? Forest Ecology and Management, 309, 4–18.

Vesterdal, L., Elberling, B., Christiansen, J. R., Callesen, I. & Schmidt, I. K. 2012. Soil respiration and rates of soil carbon turnover differ among six common European tree species. Forest Ecology and Management, 264, 185–196.

Vesterdal, L., Schmidt, I. K., Callesen, I., Nilsson, L. O. & Gundersen, P. 2008. Carbon and nitrogen in forest floor and mineral soil under six common European tree species. Forest Ecology and Management, 255, 35–48.

Wardle, D. A. & Lindahl, B. D. 2014. Ecology. Disentangling global soil fungal diversity. Science, 346, 1052–3.

White, T. J., Bruns, T., Lee, S. & Taylor, J. 1990. Amplification and Direct Sequencing of Fungal Ribosomal Rna Genes for Phylogenetics. PCR Protocols, 315–322, publisher = Academic Press.

Wickham, H. 2016. ggplot2: Elegant Graphics for Data Analysis, Springer-Verlag New York.

Xie, L., Palmroth, S., Yin, C. & Oren, R. 2024. Extramatrical mycelial biomass is mediated by fine root mass and ectomycorrhizal fungal community composition across tree species. Sci Total Environ, 950, 175175.

Yang, N., Butenschoen, O., Rana, R., Kohler, L., Hertel, D., Leuschner, C., Scheu, S., Polle, A. & Pena, R. 2019. Leaf litter species identity influences biochemical composition of ectomycorrhizal fungi. Mycorrhiza, 29, 85–96.

Zak, D. R., Pellitier, P. T., Argiroff, W., Castillo, B., James, T. Y., Nave, L. E., Averill, C., Beidler, K. V., Bhatnagar, J., Blesh, J., Classen, A. T., Craig, M., Fernandez, C. W., Gundersen, P., Johansen, R., Koide, R. T., Lilleskov, E. A., Lindahl, B. D., Nadelhoffer, K. J., Phillips, R. P. & Tunlid, A. 2019. Exploring the role of ectomycorrhizal fungi in soil carbon dynamics. New Phytol, 223, 33–39.

Zheng, H., Heděnec, P., Rousk, J., Schmidt, I. K., Peng, Y. & Vesterdal, L. 2022a. Effects of common European tree species on soil microbial resource limitation, microbial communities and soil carbon. Soil Biology and Biochemistry, 172, 108754, publisher = Pergamon.

Zheng, H., Vesterdal, L., Schmidt, I. K. & Rousk, J. 2022b. Ecoenzymatic stoichiometry can reflect microbial resource limitation, substrate quality, or both in forest soils. Soil Biology and Biochemistry, 167, 108613.

