## Supplementary material for "Tree species shape ectomycorrhizal fungal traits and community composition through litter quality and resultant soil properties": SupTable.docx

|  | Lime | | | Beech | | | Spruce | | |
| --- | --- | --- | --- | --- | --- | --- | --- | --- | --- |
|  | Odsherred | Vallø | Viemose | Odsherred | Vallø | Viemose | Odsherred | Vallø | Viemose |
| Agaricales | 0 | 0 | 2 | 1 | 0 | 0 | 0 | 0 | 1 |
| Amanita | 3 | 0 | 0 | 0 | 0 | 4 | 0 | 2 | 8 |
| Amphinema | 0 | 0 | 0 | 0 | 0 | 0 | 0 | 8 | 23 |
| Cenococcum | 1 | 0 | 4 | 1 | 0 | 1 | 4 | 1 | 0 |
| Clavulina | 0 | 0 | 0 | 0 | 0 | 0 | 0 | 0 | 1 |
| Cortinarius | 3 | 0 | 17 | 3 | 0 | 2 | 0 | 0 | 0 |
| Elaphomyces | 3 | 0 | 0 | 0 | 0 | 1 | 0 | 0 | 0 |
| Entoloma | 0 | 0 | 0 | 1 | 0 | 0 | 0 | 0 | 0 |
| Entolomataceae | 0 | 0 | 0 | 0 | 0 | 0 | 1 | 0 | 0 |
| Genea | 0 | 0 | 0 | 0 | 3 | 0 | 0 | 0 | 0 |
| Hebeloma | 1 | 1 | 0 | 1 | 0 | 0 | 0 | 0 | 0 |
| Humaria | 0 | 0 | 0 | 2 | 0 | 0 | 0 | 0 | 0 |
| Hyaloscypha | 0 | 0 | 0 | 0 | 0 | 0 | 1 | 0 | 0 |
| Hydnotrya | 1 | 0 | 0 | 0 | 7 | 2 | 0 | 0 | 0 |
| Hydnum | 0 | 0 | 1 | 0 | 0 | 0 | 0 | 0 | 0 |
| Hysterangium | 0 | 0 | 0 | 1 | 0 | 1 | 0 | 0 | 0 |
| Imleria | 0 | 0 | 0 | 0 | 0 | 0 | 6 | 2 | 0 |
| Inocybe | 0 | 2 | 1 | 0 | 0 | 0 | 0 | 0 | 0 |
| Laccaria | 0 | 13 | 3 | 2 | 4 | 0 | 0 | 0 | 0 |
| Lactarius | 0 | 0 | 0 | 10 | 15 | 6 | 0 | 2 | 0 |
| Leotia | 0 | 0 | 0 | 2 | 0 | 0 | 0 | 0 | 0 |
| Meliniomyces | 0 | 0 | 0 | 0 | 0 | 0 | 2 | 0 | 0 |
| Octaviania | 0 | 0 | 0 | 5 | 0 | 0 | 0 | 0 | 0 |
| Otidea | 0 | 0 | 7 | 1 | 0 | 0 | 0 | 4 | 0 |
| Pachyphlodes | 4 | 0 | 0 | 0 | 1 | 0 | 0 | 0 | 0 |
| Piloderma | 0 | 1 | 7 | 0 | 1 | 7 | 0 | 0 | 1 |
| Russula | 43 | 32 | 12 | 11 | 39 | 16 | 4 | 11 | 3 |
| Scleroderma | 2 | 0 | 0 | 1 | 0 | 0 | 0 | 0 | 0 |
| Sebacina | 0 | 0 | 9 | 3 | 0 | 0 | 0 | 0 | 0 |
| Thelephora | 0 | 2 | 0 | 0 | 3 | 3 | 0 | 0 | 0 |
| Thelephoraceae | 9 | 5 | 7 | 5 | 2 | 10 | 39 | 1 | 0 |
| Tomentella | 0 | 6 | 8 | 2 | 0 | 14 | 3 | 1 | 1 |
| Tuber | 3 | 1 | 0 | 1 | 0 | 0 | 0 | 0 | 0 |
| Tylopilus | 0 | 0 | 0 | 0 | 0 | 0 | 0 | 8 | 0 |
| Tylospora | 0 | 0 | 0 | 0 | 0 | 0 | 3 | 13 | 0 |
| Xerocomellus | 0 | 2 | 1 | 0 | 1 | 1 | 0 | 11 | 5 |
