## Supplementary figures and images for "Tree species shape ectomycorrhizal fungal traits and community composition through litter quality and resultant soil properties"

### Fig_S1.pdf

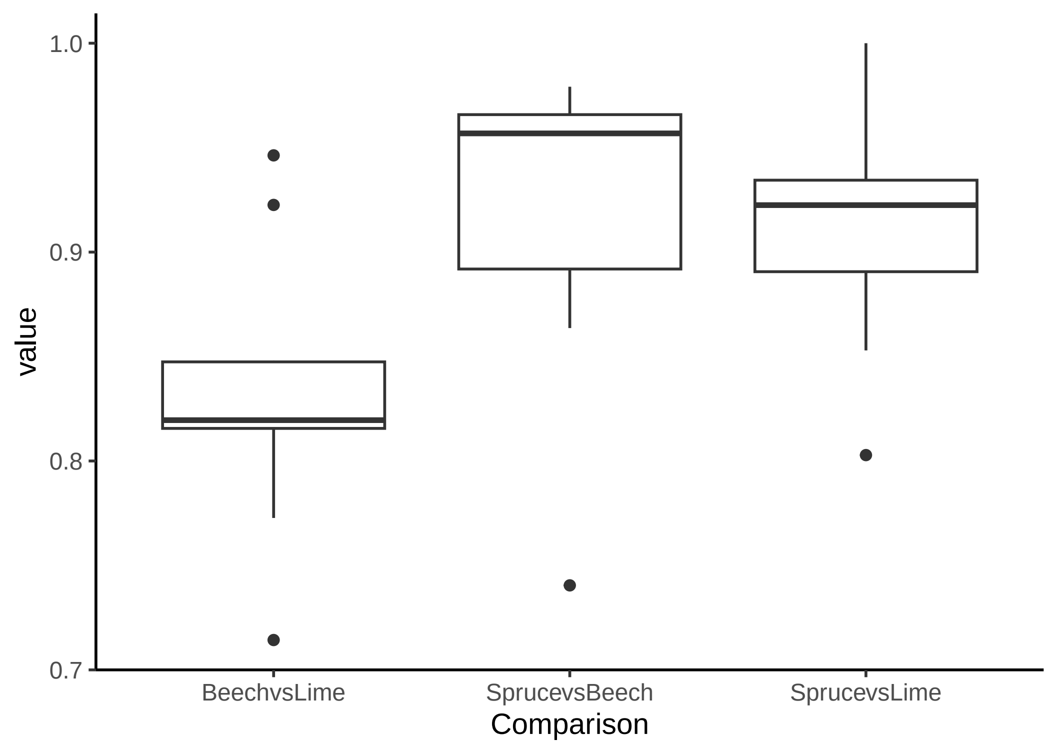
